# Rapamycin does not compromise physical performance or muscle hypertrophy after PoWeR while intermittent rapamycin alleviates glucose disruptions by frequent rapamycin

**DOI:** 10.1101/2025.03.10.642477

**Authors:** Christian J. Elliehausen, Szczepan S. Olszewski, Carolyn G. Shult, Aditya R. Ailiani, Michaela E. Trautman, Reji Babygirija, Dudley W. Lamming, Troy A. Hornberger, Dennis M. Minton, Adam R. Konopka

## Abstract

An increasing number of physically active adults are taking the mTOR inhibitor rapamycin off label with the goal of extending healthspan. However, frequent rapamycin dosing disrupts metabolic health during sedentary conditions and abates the anabolic response to exercise. Intermittent once weekly rapamycin dosing minimizes many negative metabolic side effects of frequent rapamycin in sedentary mice. However, it remains unknown how different rapamycin dosing schedules impact metabolic, physical, and skeletal muscle adaptations to voluntary exercise training. Therefore, we tested the hypothesis that intermittent rapamycin (2mg/kg; 1x/week) would avoid detrimental effects on adaptations to 8 weeks of progressive weighted wheel running (PoWeR) in adult female mice (5-month-old) by evading the sustained inhibitory effects on mTOR signaling by more frequent dosing schedules (2mg/kg; 3x/week). Frequent but not intermittent rapamycin suppressed skeletal muscle mTORC1 signaling in PoWeR trained mice. PoWeR improved maximal exercise capacity, absolute grip strength, and myofiber hypertrophy with no differences between vehicle or rapamycin treated mice. Conversely, frequent and intermittent rapamycin treated mice had impaired glucose tolerance and insulin sensitivity compared to vehicle treated mice after PoWeR; however, intermittent rapamycin reduced the impact on glucose intolerance versus frequent rapamycin. Collectively, these data in adult female mice suggest that 1) rapamycin is largely compatible with the physical and skeletal muscle benefits of PoWeR and 2) the detrimental effects of rapamycin on body composition and glucose metabolism in the context of voluntary exercise may be reduced by intermittent dosing.

## INTRODUCTION

Rapamycin (sirolimus), a mechanistic target of rapamycin (mTOR) protein kinase inhibitor, is an FDA approved drug that can extend lifespan in multiple model systems (Mannick & Lamming, 2023). Lifespan extension in UM-HET3 mice by rapamycin occurs in a dose-dependent manner and to a greater extent in females versus males (Miller et al., 2014). Rapamycin started earlier in life (9 months) may confer greater relative extension of lifespan compared to rapamycin treatment initiated in late life (20 months) (Harrison et al., 2009). In addition to lifespan, rapamycin delays several age-related pathologies, including preservation of physical and skeletal muscle function in sedentary mice and rats to combat sarcopenia and frailty (Ham et al., 2020; Joseph et al., 2019).

Despite the positive effects on lifespan and many indices of healthspan, prolonged treatment with rapamycin is associated with dose-dependent risks of metabolic side effects, including glucose intolerance, insulin resistance, and dyslipidemia (Bissler et al., 2017; Johnston, Rose, Webster, & Gill, 2008). Rapamycin acutely and potently inhibits mTOR complex 1 (mTORC1) while prolonged rapamycin treatment can inhibit mTOR complex 2 (mTORC2) signaling in culture and mice (Lamming et al., 2012; Sarbassov et al., 2006; Ye, Varamini, Lamming, Sabatini, & Baur, 2012). A leading model suggests that inhibition of mTORC1 mediates the geroprotective effects of rapamycin while off-target inhibition of mTORC2 signaling has pernicious effects on metabolic health, frailty and survival in mice (Apelo et al., 2020; Chellappa et al., 2019; Lamming et al., 2014; Mannick & Lamming, 2023; Mizunuma, Neumann-Haefelin, Moroz, Li, & Blackwell, 2014; Yu et al., 2019). Intermittent rapamycin dosing strategies more selectively inhibit mTORC1 and extend lifespan in female mice while circumventing metabolic side effects through reduced mTORC2 inhibition, (Arriola Apelo, Neuman, et al., 2016; Arriola Apelo, Pumper, Baar, Cummings, & Lamming, 2016). Due to these exciting data, an increasing number of physically active adults are now prophylactically taking rapamycin, largely using intermittent dosing schedules, even though the impact of rapamycin on the health benefits of regular physical activity and human healthspan remain unknown (Kaeberlein et al., 2023).

Preclinical and prospective studies in humans suggest engaging in either endurance and/or resistance activity is one of the most potent stimuli to protect against multi-morbidity and pre-mature mortality (Holloszy, 1997; Zhao, Veeranki, Magnussen, & Xi, 2020). Further, improvements in cardiorespiratory fitness, skeletal muscle function, and insulin sensitivity that are traditional adaptations of endurance and/or resistance exercise training are accompanied with decreased risk of morbidity and mortality (Abou Sawan, Nunes, Lim, McKendry, & Phillips, 2023; Imboden et al., 2018; Lanza et al., 2008).

To model adaptations commonly observed after combined endurance and resistance exercise in humans, we and others have recently implemented progressive weighted wheel running (PoWeR) in adult and aged mice (Dungan et al., 2022, 2019; Englund et al., 2020; Elliehausen et al. 2025). PoWeR is a voluntary, high-volume exercise paradigm that increases muscle fiber size and oxidative capacity in multiple muscles while improving whole body glucose metabolism and physical function. We have previously demonstrated that short-term PoWeR in adult mice stimulates mTORC1 signaling as evident by a 2-3 fold increase in the phosphorylation of downstream substrates p70S6K (T389) and rpS6 (S233/356) particularly in muscles that increase in mass after longer-term PoWeR (Elliehausen et al. 2025). mTORC1 signaling is a critical regulator of skeletal muscle anabolism whereby disrupting mTORC1 signaling genetically or by rapamycin attenuates the muscle protein synthetic and hypertrophic response to exercise in rodents and humans (Bodine et al., 2001; Drummond et al., 2009; Goodman et al., 2011; Gundermann et al., 2014; Ogasawara et al., 2016; Philp et al., 2015; You et al., 2019). Similarly, genetic knockouts of *Rictor*, a core component of mTORC2, reveal blunted exercise induced glucose uptake and muscle protein synthesis (Kleinert et al., 2017; Ogasawara, Knudsen, Li, Ato, & Jensen, 2020). Improved functional adaptations to muscle mass, hypertrophy, insulin sensitivity, and physical performance are associated with repeated increased signaling through mTORC1/2 (Egan & Sharples, 2022).

Collectively, these data support the current status quo that rapamycin-mediated inhibition of mTORC1/2 is largely contraindicated with exercise. However, the majority of preclinical studies to date have only provided rapamycin frequently at high doses or on the same day as the exercise stimuli and have primarily evaluated the effects on the hypertrophic and protein synthetic responses to models of simulated resistance or acute forced treadmill exercise (Goodman et al., 2011; Ogasawara et al., 2016; Philp et al., 2015). It is also unknown whether inhibition of mTORC1 by rapamycin during a physiological, voluntary, and high-volume model of exercise impairs systemic and skeletal muscle adaptations and whether these effects vary with different rapamycin dosing schedules.

Therefore, the goals of this study were to 1) identify if mTOR inhibition via rapamycin will alter the physical, metabolic, and skeletal muscle adaptations to PoWeR in adult female mice and 2) determine whether there are dosing regimen-dependent outcomes that would support or refute the concomitant use of rapamycin and exercise in future healthspan and lifespan studies. Female mice were *a priori* selected for the current study because 1) the lifespan extending effects of rapamycin are relatively greater in females than males and 2) female mice run greater volumes and more consistently than males (Konhilas et al., 2004; Mannick & Lamming, 2023; Miller et al., 2014). Here, we demonstrate that frequent (3x/wk) and intermittent (1x/wk) rapamycin dosing is compatible with the physical performance and muscle hypertrophy benefits of PoWeR. However, we identify frequent rapamycin disrupts whole-body glucose homeostasis during PoWeR which is reduced by intermittent dosing. These results provide support for concomitant rapamycin and exercise interventions aimed at evaluating lifespan and healthspan, although alternative dosing regimens may be warranted to minimize metabolic disruptions.

## RESULTS

### Influence of Rapamycin on Running Behavior and Body Composition after PoWeR

We first sought to determine if different dosing schedules of rapamycin altered spontaneous running behavior and body composition during PoWeR. At baseline, there were no differences in running volume, body weight or body composition between groups. Quantification of cumulative running volume revealed mice treated with intermittent rapamycin (Rapa 1x/wk) ran more over the course of the training intervention compared to the other treatment groups (**Fig. 1A & B**) despite receiving the same number of total i.p injections. Weekly assessment of bodyweight did not differ between groups over the course of the intervention (**Fig. 1C**), however, PoWeR trained mice treated with vehicle or intermittent rapamycin reduced adiposity while mice treated with frequent rapamycin did not (**Fig. 1D & E**).

**Figure 1:**
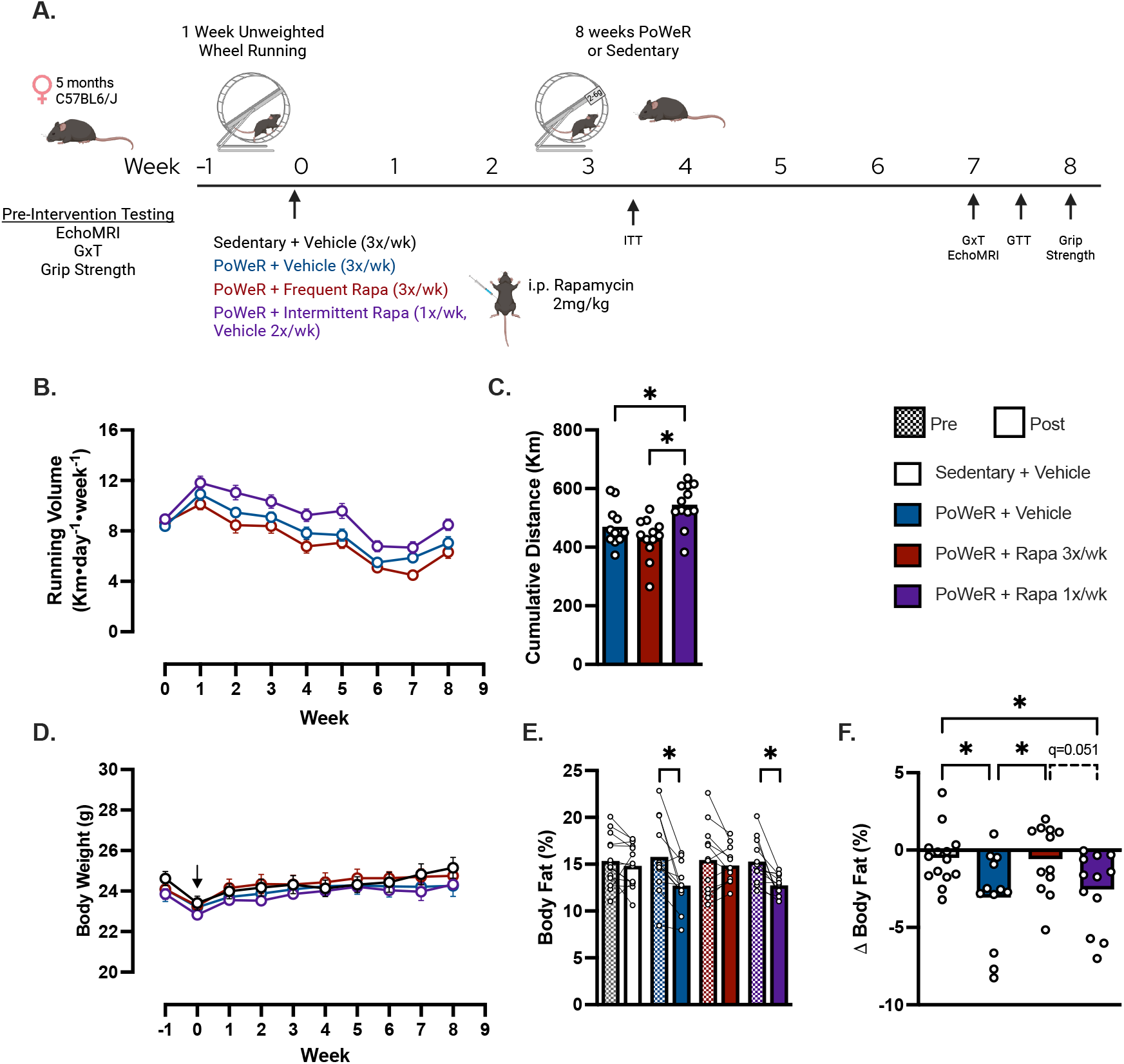
Physiological Characterization. **(A)** Study schematic and timepoints for key measurements. **(B)** Average daily voluntary running distance per week and **(C)** cumulative volume over 1 week of unweighted wheel running and 8 weeks of PoWeR. **(D)** Weekly bodyweight. **(E)** Body fat percentage pre and post intervention. **(F)** Delta (post minus pre) body fat percentage. Arrow denotes randomization of wheel running mice and the start of vehicle and rapamycin treatment. N=11-14 per group. Data presented as mean plus individual data points or error bars represent SEM. Data analyzed by 2-way ANOVA with repeated measures was used to assess main effects for time, treatment, and time x treatment interactions and multiple post hoc comparisons were FDR corrected with a two-stage step-up method of Benjamini, Krieger, and Yekutieli (E) otherwise one-way ANOVA with multiple comparisons FDR corrected by a 2-stage step-up method (C, F). *q<0.05. Panel A created in BioRender.com. GxT-graded exercise test; ITT-insulin tolerance test; GTT-glucose tolerance test.

### Disruptions to glucose metabolism with frequent rapamycin during PoWeR are reduced by intermittent rapamycin

Impaired glucose homeostasis is a negative metabolic side effect of frequent administration of rapamycin or rapalogs in sedentary humans (Bissler et al., 2017; Johnston et al., 2008). In mice, disruptions to glucose metabolism can be observed as early as 2 weeks of rapamycin treatment (Lamming et al., 2012; Yang et al., 2012). Therefore, to determine the impact of rapamycin on glucose metabolism after PoWeR, we evaluated fasting glucose, glucose tolerance, and insulin tolerance after 3- or 7-weeks following rapamycin treatment onset (**Fig. 1A**). All tolerance tests were performed 24-hr after the last exercise bout and 24-hr or 72-hr after the last frequent and intermittent rapamycin dose, respectively. 72-hr was selected for the intermittent group as this was suggested to be when mTORC1 inhibition is alleviated in skeletal muscle in sedentary mice receiving intermittent rapamycin (2mg/kg, 1x/wk) (Arriola Apelo, Neuman, et al., 2016)

Following an overnight fast (∼16-hr) PoWeR trained mice treated with vehicle had lower (q=0.05) fasting blood glucose compared to sedentary control (**Fig. 2A**). PoWeR trained mice treated with frequent rapamycin had greater fasting blood glucose compared to PoWeR + vehicle and intermittent rapamycin and was not different from sedentary mice (**Fig. 2A**). Frequent and intermittent rapamycin both disrupted glucose tolerance after PoWeR compared to vehicle control as evident by a 40% and 21% greater glucose burden determined by the area of the blood glucose curve (**Fig. 2B**). However, glucose intolerance was less severe in intermittent versus frequent rapamycin treated mice after PoWeR (**Fig. 2B**).

**Figure 2:**
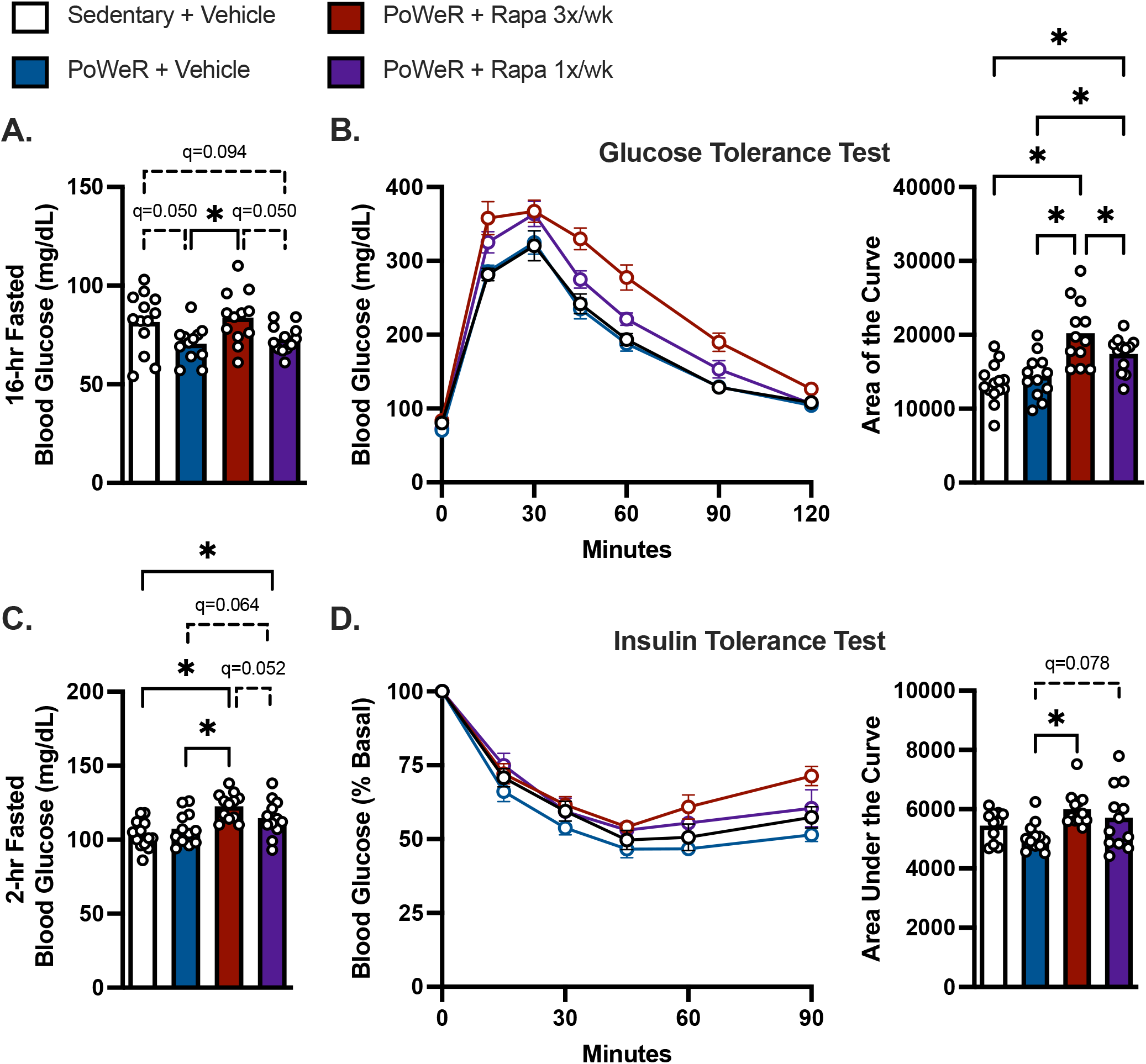
Disruptions to glucose homeostasis by frequent rapamycin are less severe by once weekly intermittent dosing during PoWeR. **(A)** Overnight fasted (∼16-hrs) blood glucose level prior to glucose injection. **(B)** Blood glucose tolerance and area of the curve calculated as the area above baseline starting blood glucose value prior to glucose injection. **(C)** Fasted (2-hrs) blood glucose prior to insulin injection. **(D)** Blood glucose curve during an insulin tolerance test and area under the curve. N=11-14 per group. Data presented as mean plus individual data points or error bars represent SEM. Data analyzed by one-way ANOVA with multiple comparisons FDR corrected by a 2-stage step-up method (A-D). *q<0.05

Following a 2-hr fast and prior to the insulin tolerance test, frequent and intermittent rapamycin treatments resulted in elevated fasting glucose levels compared to sedentary controls (**Fig. 2C**). After PoWeR, mice treated with frequent but not intermittent rapamycin (q=0.078) were relatively less insulin sensitive compared to vehicle treated mice determined by the area under the relative (% Basal) blood glucose curve (**Fig. 2D**). Collectively, these data suggest the disruptions to glucose tolerance with frequent rapamycin treatment are not overcome with exercise training. Further, we identify glucose intolerance is less severe with intermittent rapamycin compared to frequent rapamycin during PoWeR.

### Rapamycin does not attenuate the improvement in whole-body physical capacity nor myofiber hypertrophy after PoWeR

We next aimed to determine if rapamycin would alter the physical performance benefits and skeletal muscle hypertrophy from PoWeR. Despite differences in running volume and body composition after PoWeR between vehicle, intermittent and/or frequent rapamycin treated mice, rapamycin did not influence the increase in maximal running capacity nor absolute all-limb grip strength after PoWeR (**Fig. 3A-D**).

**Figure 3:**
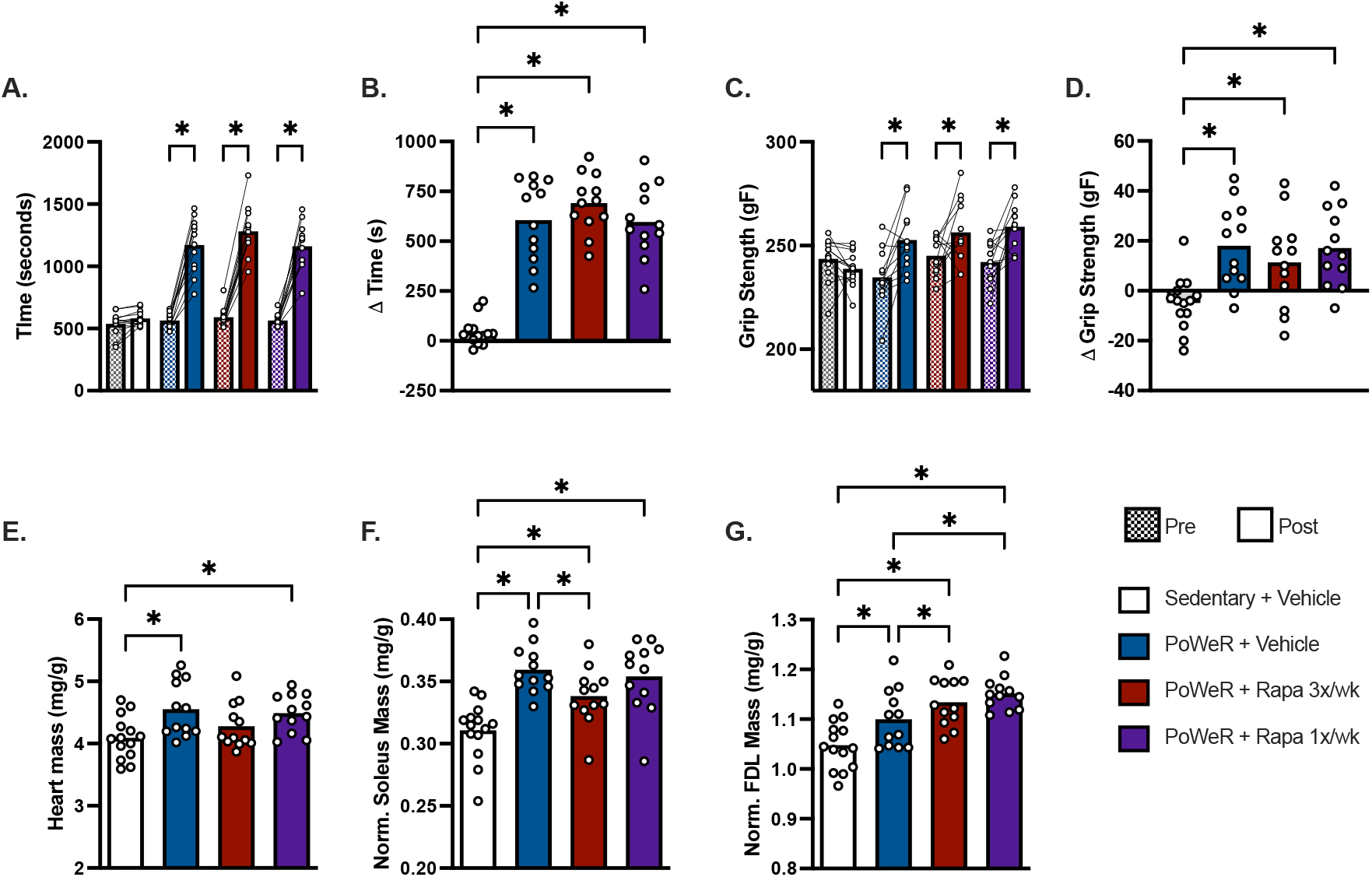
Rapamycin does not impair physical performance adaptions to PoWeR. **(A)** Time run during a graded exercise test. **(B)** Delta (post minus pre) time run. **(C)** All-Limb grip strength. **(D)** Delta (post minus pre) all-limb grip strength **(E)** Heart mass normalized to bodyweight. **(F)** Soleus muscle mass normalized to bodyweight. **(G)** FDL muscle mass normalized to bodyweight. N=12-14/group. Data presented mean plus individual data points. Data analyzed by 2-way ANOVA with repeated measures was used to assess main effects for time, treatment, and time x treatment interactions and multiple post hoc comparisons were FDR corrected with a two-stage step-up method of Benjamini, Krieger, and Yekutieli (A & C) otherwise one-way ANOVA with multiple comparisons FDR corrected by a 2-stage step-up method (B, D-G). *q<0.05.

Considering the notion that inhibition of mTORC1 by rapamycin would blunt muscle hypertrophy, we evaluated mass and/or fiber type specific cross sectional area (CSA) in the heart, soleus and FDL which we and others have identified to respond to PoWeR (Dungan et al., 2022, 2019; Englund et al., 2020; Elliehausen et al., 2025). Frequent but not intermittent rapamycin attenuated the increase in heart mass normalized to body mass after PoWeR (**Fig 3C**), although the impact on heart mass by frequent rapamycin did not seem to significantly impact maximal exercise performance. PoWeR increased soleus and FDL muscle mass normalized to body mass in vehicle, frequent, and intermittent rapamycin treated mice (**Fig 3D & E**). However, we identified muscle specific effects of rapamycin on muscle mass gains after PoWeR. In the oxidative soleus, frequent but not intermittent rapamycin treated mice had lower soleus muscle mass after PoWeR compared to vehicle control (**Fig 3D**). Conversely, intermittent and frequent rapamycin had greater FDL muscle mass after PoWeR compared to vehicle control (**Fig 3E**).

Performing immunohistochemistry on whole muscle cross sections permitted the evaluation of fiber-type specific CSA in the soleus and FDL (**Fig 4A & B**). In the soleus, PoWeR trained mice treated with intermittent rapamycin had ∼40% greater type IIA CSA compared to sedentary controls while PoWeR trained mice treated with vehicle (+24%, q=0.07) or frequent rapamycin (+20%, q=0.1) had non-significantly greater Type IIA CSA compared to sedentary (**Fig 4C**). In the FDL, PoWeR trained mice had greater Type I and IIA myofiber CSA compared to sedentary, with no differences detected between PoWeR trained groups (**Fig 4D**).

**Figure 4.**
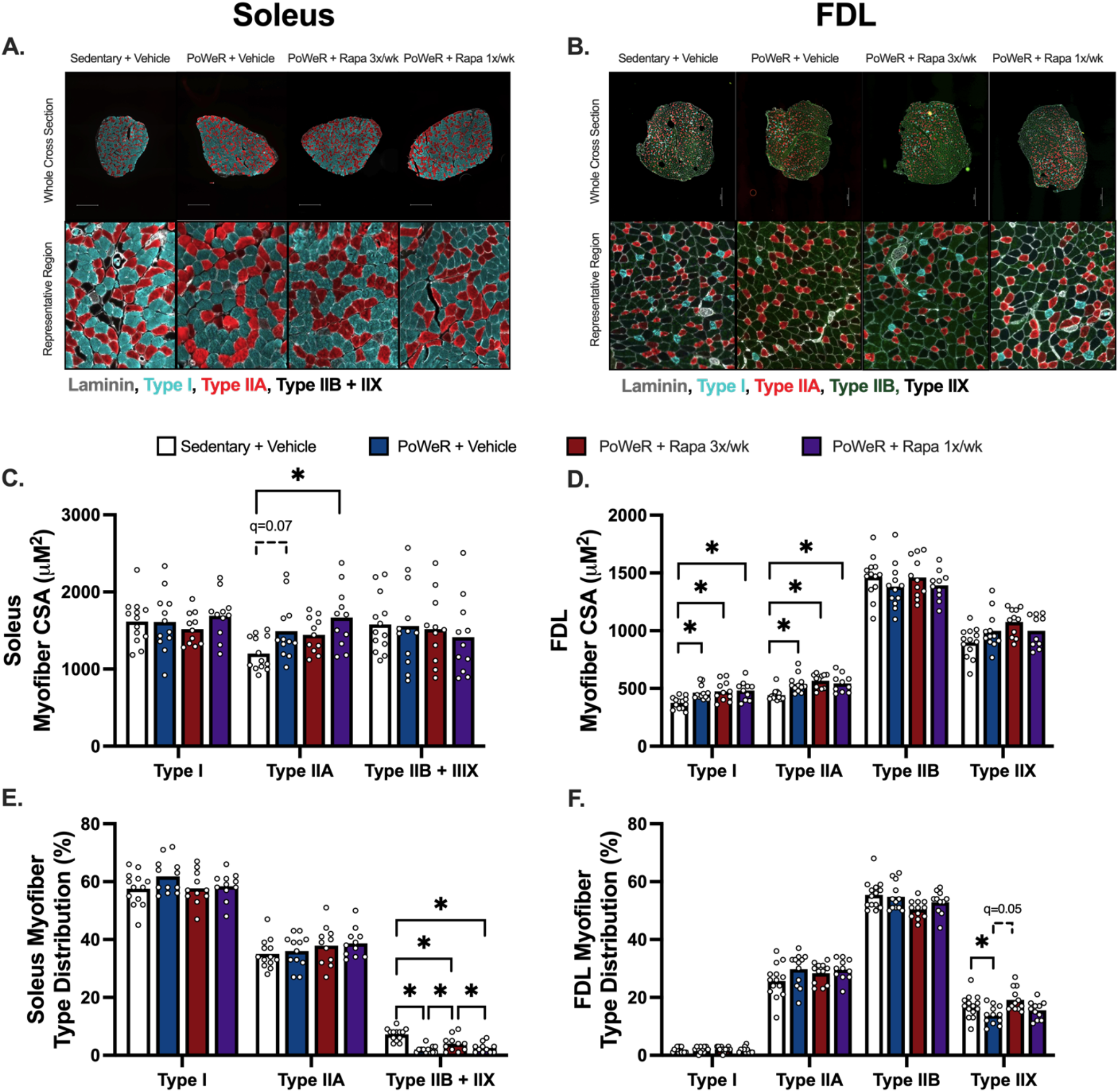
Myofiber hypertrophy in the soleus and FDL after PoWeR are not impeded by rapamycin. Representative mid-belly cross sections of the **(A)** soleus and **(B)** FDL subjected to immunohistochemistry for myofiber typing (Type I, Type IIA, Type IIB and Type IIX.) Due to low abundances of IIB and IIX myofibers in the soleus these fibers were pooled. Average myofiber size per myofiber type in the **(C)** soleus and **(D)** FDL. Myofiber type distribution in the **(E)** soleus and **(F)** FDL. N=11-13 per group. Data presented as mean plus individual data points. Each myofiber type was analyzed by one-way ANOVA with multiple comparisons FDR corrected by a 2-stage step-up method (C-F). *q<0.05. scale bar in A, B = 500μM

All PoWeR trained mice, independent of rapamycin treatment, had a lower proportion of type IIB+IIX myofibers in the soleus compared to sedentary (**Fig 4E**). However, the proportion of IIB+IIX myofibers was greater with frequent rapamycin than intermittent rapamycin and vehicle in PoWeR trained mice. Myofiber type distribution in the FDL was largely unaffected by PoWeR with or without rapamycin treatment, with only the frequent rapamycin group having a small but statistically significant greater proportion of FDL Type IIX fibers compared to vehicle (**Fig 4F**). These results confirm PoWeR increases Type I and/or IIA fiber size while rapamycin did not negatively impact myofiber hypertrophy after PoWeR. These data also highlight a discrepancy between muscle mass and myofiber size urging caution when interpreting mass increases as muscle hypertrophy.

### Skeletal muscle mTOR signaling

Finally, we aimed to evaluate mTORC1 and mTORC2 signaling in multiple skeletal muscles to associate with the observed physiological and skeletal muscle outcomes. Skeletal muscles were collected following a 24-hr running wheel lock and 2-hr fast. The last dose of frequent rapamycin was provided 24-hr prior to tissue collection while the intermittent rapamycin was dosed 7-days prior. We evaluated phosphorylation of ribosomal protein S6 (rpS6 S235/236), a downstream target of mTORC1 signaling, in the tibialis anterior (TA), soleus, and FDL. In the TA, greater phosphorylation of rpS6 S235/236 was observed in PoWeR trained mice treated with vehicle or intermittent rapamycin. Conversely, phosphorylation of rpS6 S235/236 was no different between PoWeR trained mice treated with frequent rapamycin versus sedentary controls suggesting frequent rapamycin suppressed rpS6 S235/236 signaling with PoWeR (**Fig 5A**). In the FDL, there was no increase in the phosphorylation of rpS6 S235/236 in PoWeR trained mice treated with vehicle or intermittent rapamycin (**Fig 5B**), however, frequent rapamycin suppressed the phosphorylation of rpS6 S235/236 compared to both PoWeR trained vehicle and intermittent rapamycin groups as well as sedentary controls. In the soleus muscle, we found all PoWeR trained groups had lower phosphorylation of rpS6 S235/236 compared to sedentary controls (**Fig 5C**).

**Figure 5:**
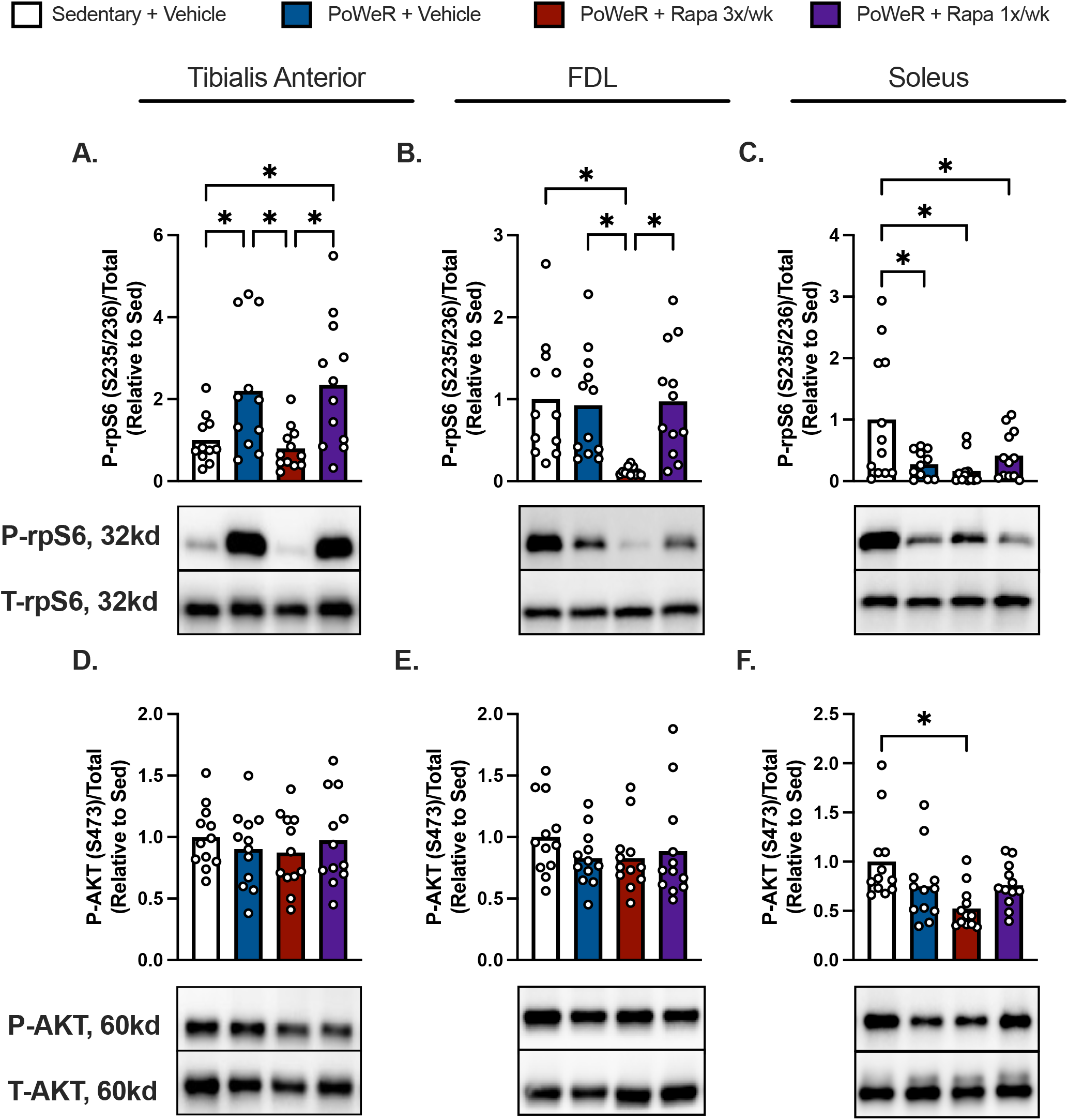
Skeletal muscle mTOR signaling following PoWeR training. Ratio of phosphorylated rpS6 (S235/236) to total protein via immunoblotting in the **(A)** Tibialis Anterior, **(B)** FDL, and **(C)** Soleus with reflective immunoblot panels below corresponding to the treatment group order in the figures. Ratio of phosphorylated AKT (S473) to total protein via immunoblotting in the **(D)** Tibialis Anterior, **(E)** FDL, and **(F)** Soleus. N=11-12 per group. Data presented as mean plus individual data points. Data analyzed by one-way ANOVA with multiple comparisons FDR corrected by a 2-stage step-up method (A-F). *q<0.05.

Since disruption to mTORC2 signaling is a proposed mechanism for metabolic dysfunction with prolonged rapamycin treatment we probed for phosphorylated AKT S473 as a substrate of mTORC2. We identified no differences in the phosphorylation of AKT S473 between any groups in the TA or FDL (**Fig 5D & E**). However, in the soleus we did detect lower AKT S473 in PoWeR trained mice treated with frequent rapamycin compared to sedentary controls (**Fig 5F**).

## DISCUSSION

While exercise and rapamycin can independently improve multiple biological and physiological indices of aging, it has been speculated that rapamycin and exercise are incompatible due to their opposing effects on mTOR signaling (Elliehausen et al., 2023). To address this fundamental question, the primary goals of this study were to 1) determine the influence of rapamycin on mTOR signaling and cellular and physiological adaptations to PoWeR and 2) whether these responses differed depending on dosing schedule in adult female mice. The major findings from this study were that despite differences in dosing frequency, neither frequent nor intermittent rapamycin undermined the improvement in physical capacity and myofiber hypertrophy by PoWeR. Frequent and intermittent rapamycin impaired glucose tolerance during PoWeR however, the detrimental effects of rapamycin on glucose homeostasis were reduced by once weekly intermittent dosing. These data suggest that rapamycin is largely compatible with the majority of the health benefits of PoWeR and that intermittent once weekly dosing may be an approach to partially mitigate negative metabolic side effects of rapamycin during PoWeR in adult female mice.

### Metabolic Function

Frequent administration of rapamycin at doses consistent with the FDA label increase risk for glucose intolerance and insulin resistance, which are known risk factors for cardiometabolic disease and would appear counter to healthy aging (Bissler et al., 2017; Johnston et al., 2008). Further, inhibition of mTORC1 and/or mTORC2 signaling attenuate the acute increase in muscle protein synthesis and glucose uptake after a single bout of exercise and diminishes muscle hypertrophy after models of rodent resistance exercise (Goodman et al., 2011; Ogasawara et al., 2016, 2020; Ogasawara & Suginohara, 2018). Despite these undesirable side effects, a growing number of people, particularly younger to middle aged, are prophylactically consuming rapamycin off label with the intent to delay age-associated conditions, in lieu of several ongoing human clinical trials are testing if different rapamycin dosing strategies can safely and effectively improve the biology and physiology of aging and age-associated diseases (Konopka & Lamming, 2023). However, it remained unknown if rapamycin would alter the multitude of health benefits, like glycemic control, from voluntary exercise.

Here we show that despite engaging in a high-volume exercise paradigm, mice treated with frequent rapamycin were glucose intolerant and insulin resistant compared to vehicle control.

Importantly, intermittent once weekly dosing reduced the severity of impairments to whole-body glucose homeostasis by frequent rapamycin after PoWeR but did not completely protect against glucose intolerance relative to vehicle control. Previous reports identified intermittent dosing once every 5 or 7 days prevents glucose intolerance and reduces pyruvate intolerance observed with more frequent rapamycin dosing (Arriola Apelo, Neuman, et al., 2016; Arriola Apelo, Pumper, et al., 2016). While our findings are largely in agreement with previous findings in sedentary mice whereby intermittent rapamycin reduced metabolic disruptions observed with frequent rapamycin, intermittent rapamycin still imparted some glucose intolerance relative to PoWeR trained mice treated with vehicle. An important distinction between current study with PoWeR and the previous work in sedentary mice is the time of glucose tolerance testing relative to the last dose of rapamycin. Here we performed glucose tolerance testing 3 days after the last dose while the prior studies completed glucose tolerance testing 5 or 7 days after the last dose (Arriola Apelo, Neuman, et al., 2016). Regardless, the findings from this study are in line with the notion that intermittent dosing strategies may be effective at mitigating glucose disruptions by more frequent rapamycin even in the context of PoWeR.

### Physiological Adaptations

Our data indicate that rapamycin does not impede engagement in voluntary exercise or impact the increase in exercise capacity after PoWeR. These results are largely supported by previous findings where a single dose of rapamycin did not impact the increase in mitochondrial protein synthesis after acute treadmill exercise, and frequent rapamycin treatment did not compromise exercise capacity nor spontaneous physical activity in sedentary mice (Ham et al., 2020; Philp et al., 2015; Ye et al., 2013). Cardiorespiratory fitness or VO_2_ max is one of the strongest predictors of morbidity and mortality in humans (Imboden et al., 2018; Mandsager et al., 2018). While we measured treadmill exercise capacity as a surrogate for cardiorespiratory fitness, the lack of impact of rapamycin on the improvement in exercise capacity after PoWeR is promising for the goal of reducing risk of all-cause mortality and improving healthy longevity.

Interestingly, frequent rapamycin appeared to prevent the decline in adiposity observed with PoWeR in the vehicle and intermittent rapamycin mice. These results are paradoxical to the literature, where previous evidence suggests rapamycin decreased adiposity, particularly in diet induced obesity models (Houde et al., 2010). Alternatively, models of mTORC1 activation also reduce adiposity (Magdalon et al., 2016). There may be a fine-tuned activity of mTORC1/2 signaling necessary to regulate adiposity in the context of exercise training that frequent rapamycin treatment may impair, though this remains to be elucidated. However, since mice treated with intermittent rapamycin ran a greater cumulative distance, further studies are needed to determine if the loss of adiposity as well as the reduced impact on glucose intolerance are due to direct effects of rapamycin dosing schedule or are secondary to differences in running volume between intermittent and frequent rapamycin treatment.

### Muscle Mass, Size, and Grip Strength

We hypothesized frequent rapamycin treatment would mitigate the increase in skeletal muscle mTORC1 signaling, mass, hypertrophy, and grip strength after PoWeR. This hypothesis was supported by previous findings by our lab and others which have shown that PoWeR induced mTORC1 signaling, muscle hypertrophy, and improved whole-body physical performance (Dungan et al., 2022, 2019; Englund et al., 2020; Elliehausen et al. 2025) while inhibition of mTORC1 signaling by high doses of rapamycin attenuated the muscle protein synthesis response to a single bout of resistance exercise in humans (Drummond et al., 2009; Gundermann et al., 2014) and muscle hypertrophy after non-voluntary models of resistance exercise in rodents (Goodman et al., 2011; Ogasawara et al., 2016). We probed for rpS6 in muscles after euthanasia to gather insight into the mTORC1 inhibition predicted with our dosing regimens. After a 24-hr washout from exercise and the last frequent rapamycin dose we observed marked suppression of downstream mTORC1 signaling in multiple muscles which we interpret to suggest that 3x/wk dosing regimen was sufficient to repeatedly inhibit mTORC1 over the course of the training intervention, at least 24-hr after the last dose in the context of our study. In contrast we presume intermittent dosing only suppressed mTORC1 signaling acutely between doses considering readouts of mTORC1 signaling were equivalent to vehicle controls by the 7^th^ day after dosing. While frequent rapamycin appeared to attenuate the increase in soleus mass after PoWeR, this was not detected at the level of the myofiber. In the context of our frequent dosing regimen, the mTORC1 inhibition by rapamycin did not compromise myofiber hypertrophy in the soleus or FDL, nor the improvement in whole-limb grip strength after PoWeR. Therefore, it is possible the residual amount of mTORC1 signaling detected after PoWeR in the frequent rapamycin treated mice is sufficient to support muscle growth. Alternatively, rapamycin sensitive mTORC1 signaling may be dispensable for the benefits of PoWeR. Conversely, alternative pathways may compensate for mTORC1 inhibition or may serve as the primary drivers for PoWeR-induced muscle hypertrophy as evident by recent findings that mTORC2 signaling, the mitogen activated protein kinase (MAPK) pathway, or the transcription factor MYC are also mechanisms regulating skeletal muscle hypertrophy (Murach et al., 2022; Ogasawara, Jensen, Goodman, & Hornberger, 2019; Steinert et al., 2021).

### Limitations/Considerations

In the current study, we focused on female mice which are traditionally underrepresented in exercise research. While additional studies to compare sex-differences will be needed, we chose to first study female mice because they have more consistent running behavior during voluntary wheel running and relatively greater lifespan extension by rapamycin compared to males (Konhilas et al., 2004; Mannick & Lamming, 2023; Miller et al., 2014). Further, we successfully randomized mice to match for running volume as shown by equivalent running volumes in each group during the first week. Despite this control, PoWeR trained mice receiving intermittent rapamycin accumulated greater running volume than mice treated with vehicle or frequent rapamycin. While this is a significant finding that supports the concept that intermittent rapamycin may enhance spontaneous running behavior, the greater running volume could contribute to intermittent rapamycin mitigating the negative effects of frequent rapamycin on fasted blood glucose, glucose intolerance, and adiposity after PoWeR. However, we did not identify associations with total running volume and any outcome differentially impacted by intermittent versus frequent rapamycin (data not shown), suggesting the divergent effects are likely due to rapamycin dosing schedule. However, future studies are needed to distinguish the direct impact of the dosing schedule versus training volume. Further, it will be important to validate if greater running volume may be a benefit of intermittent rapamycin dosing schedules since greater physical activity levels are associated with lower risk of morbidity and mortality in humans (Zhao et al., 2020). Last, this study did not include a sedentary group of mice treated with intermittent or frequent rapamycin. Omission of sedentary treatment groups limits the ability to detect the independent effects of rapamycin versus PoWeR on physiological and skeletal muscle outcomes per se. However, a goal of this study was to identify if the adaptations of PoWeR were impeded by the addition of various rapamycin doses regimens, and we contextualize these observations by leveraging data from seminal studies highlighting the effects of rapamycin in sedentary mice when available.

## Conclusion

mTORC1 is considered a central regulator of exercise adaptions. In preclinical and human clinical studies, inhibition or ablation of mTORC1 reduced muscle protein synthesis and muscle hypertrophy. Therefore, the dogma was that rapamycin would not be compatible with the health benefits of exercise despite never being formally tested. Contrary to our hypothesis, the data from the current study suggest rapamycin treatment is largely compatible with the physical performance benefits of PoWeR, however, more frequent dosing may attenuate the improvement in glucose homeostasis and loss of adiposity after PoWeR. Intermittent rapamycin treatment concomitant with PoWeR mitigated the majority of the negative effects of more frequent rapamycin treatment, suggesting the potential for intermittent rapamycin dosing schedules to be a viable strategy to test under long-term or lifelong trials for healthy longevity and lifespan.

## Methods

### Animal Use and Care/Ethical Approval

All animal procedures were performed in conformance with institutional guidelines and were approved by the Institutional Animal Care and Use Committee of the William S. Middleton Memorial Veterans Hospital. Female C57BL/6J were procured from the Jackson Laboratory (000664) at 18 weeks of age. Mice were acclimated to the animal research facility for 2-weeks before entering studies. All mice were singly housed provided Purina Rodent Chow (5001) *ad libitum* and provided enrichment to minimize stress. The animal room was maintained with a 12:12 h light-dark cycle at 23°C.

### PoWeR Training

Progressive weighted wheel running (PoWeR) was used as previously published (Dungan et al., 2019). Briefly, 20-week-old female C56BL6/J mice (n=50) were acclimated to unweighted running wheels for 7 days. Mice that did not engage in 1 week of unweighted wheel running were allocated to the sedentary group (sedentary + vehicle control (n=14, 3x/wk). The remaining 36 mice were randomized to the following groups to match for running volume and body composition: PoWeR + vehicle (n=12, 3x/wk), PoWeR + frequent rapamycin (3x/wk; n=12), PoWeR + intermittent Rapamycin (1x/wk, n=12). After, one week of unweighted wheel running, sedentary mice did not have access to a running wheel and housed in cages of similar size with a hut and bedding for enrichment. For PoWeR, 1-gram magnets were fixed to one side of an 11cm diameter wheel to asymmetrically load. Weights were progressively added starting at 2g until 6g at week 6 as previously performed (Dungan et al., 2019) for a total of 8-weeks of weighted wheel running. Weekly average of daily wheel running distance (km/day) was measured by ClockLab Software (Actimetrics, Wilmette, IL).

### Rapamycin Treatment

Rapamycin (LC Laboratories) was dissolved in ethanol and stored at -80°C. On injection days rapamycin was thawed and mixed with filter sterilized solution of 5% PEG 400 and 5% Tween-80 in 0.9% saline. Vehicle treated mice received equal parts ethanol without rapamycin in 5% PEG 400 and 5% Tween-80 in 0.9% saline. Injection volume ranged between 80-120uL with the intended rapamycin concentration at 2mg/kg bodyweight. All mice received equal handling and injections which occurred 3 days per week (Monday/Wednesday/Friday) between the hours of 8am-10am. Mice treated with rapamycin 1x/wk received rapamycin on Monday and vehicle Wednesday and Friday.

### Body Composition

Body composition using an EchoMRI 3-in-1 body composition analyzer was completed on all mice before and after the 7^th^ week of PoWeR. Mice were placed into a restraint tube (no anesthesia) for <2 min of total time generating duplicate measures of fat free mass (FFM) and fat mass. Duplicate measures were averaged for final data use.

### Graded exercise test

Graded exercise tests were performed on a Columbus 3/6 treadmill (Columbus Instruments, Columbus OH) before (Pre) and after 8 weeks of PoWeR. Mice were acclimated for 2 consecutive days prior to testing. Acclimation consisted of mice sitting on a stationary treadmill for 3 minutes with the shock grid activated to (3hz and 1.5mA). Next, the treadmill was inclined to 10 degrees and speed of 6m/min for 5 minutes then progressively increased to 12m/min for 5 more minutes for a total of 10 minutes of treadmill movement. A graded exercise test to exhaustion was performed following a previously published method for measuring VO_2_ max in mice (Petrosino et al., 2016). Exhaustion was defined as spending 5 consecutive seconds in contact or repeated contact with the shock grid demonstrating an inability to reengage treadmill running. For post-training testing, mice completed one acclimation day prior to graded exercise testing. All acclimations and graded exercise tests were performed in the early dark phase approximately 30-90 minutes after light-dark transition. Investigator was blinded to treatment groups during testing.

### Insulin and Glucose Tolerance Tests

Insulin tolerance tests (ITT) were performed during the 4^th^ week of PoWeR, 24 hours after the last exercise bout and 24 or 72 hours after the previous rapamycin dose for the frequent (3x/wk) and intermittent (1x/wk) rapamycin groups respectively. Following a 2-hour daytime fast beginning at 8:00am, mice were i.p. injected with insulin (0.75 U/kg bodyweight). Blood glucose was recorded from a tail vein bleed before (0 minutes) and 15, 30, 45, 60 and 90 minutes post injection. Investigator was blinded to treatment groups during testing.

The glucose tolerance test (GTT) was performed during the 8^th^ week of PoWeR, 24 hours after the previous exercise bout and 24 or 72 hours after the previous rapamycin dose for the frequent (3x/wk) and intermittent (1x/wk) rapamycin groups respectively Following an overnight fast (∼18hr), mice were i.p. injected with glucose (2mg/kg bodyweight). Blood glucose was recorded from a tail vein bleed before (0 minutes) and 15, 30, 45, 60, 90, and 120 minutes post injection. Blood glucose was measured using a Bayer Contour Blood glucose meter and glucose strips. Investigator was blinded to treatment groups during testing.

### Grip Strength

Grip strength was evaluated before and after the 8^th^ week of PoWeR by measuring peak force production of all limbs (fore and hind) using a Grip Strength Meter (Columbus Instruments, Columbus, OH). Mice were placed on the horizontal grid connected to a force transducer. Mice were held by the base of the tail and when the mouse fully gripped the grid, the mouse was pulled horizontally at a consistent speed until the grip released. This test was repeated 3 times with 15 minutes between tests. Grip strength was measured as highest grams force across the 3 attempts. Testing was performed by a blinded investigator in the early dark phase to align with natural circadian rhythms. Investigator was blinded to treatment groups during testing.

### Animal Dissections

Running wheels were locked for 24 hours prior to euthanasia. Rapamycin was dosed 24 hours prior to euthanasia in the frequent rapamycin group (3x/wk) and 7 days prior to euthanasia in the intermittent group (1x/wk) to represent the steady state conditions the mice would be experiencing during the intervention prior to the next dose. Euthanasia was performed via cervical dislocation following a sub mandibular puncture for blood collection in the morning following a ∼2-3 hour fast. Muscles from both legs (tibialis anterior (TA), soleus, and flexor digitorum longus (FDL)) and heart were rapidly excised, weighed, and prepared for future biochemical analyses as outlined in each section. Muscle weights were expressed relative to bodyweight. Each tissue was dissected by the same investigator to limit variability.

### Muscle Immunohistochemistry

Soleus and FDL from the left leg were weighed, measured for muscle length, and submerged in optimum cutting temperature compound (OCT, Tissue-Tek; Sakura Finetek, The Netherlands) at resting length and frozen in liquid nitrogen chilled isopentane as previously performed (Trautman et al., 2023).

### Immunoblotting

The tibialis anterior, FDL, and soleus were powdered, prepared, and immunoblotted blotted as previously detailed (Elliehausen et al 2025). Antibodies used were P-rpS6 (S235/236) (#4858), rpS6 (#2217), P-AKT (S473) (#4060), AKT (#4961) all obtained from Cell Signaling Technologies.

### Statistical Analysis

Data in bar graphs presented as mean with individual data points. Data in line graphs are presented as mean with error bars expressed as standard error of the mean. One-way ANOVA was used to compare the effects between Sedentary + Vehicle and PoWeR + Vehicle, PoWeR + Rapa 3x/wk, and PoWeR + Rapa 1x/wk. For comparisons with pre- and post-measures, groups were compared with a two-way ANOVA with repeated measures to determine the main effects of 1) Time or 2) Treatment. Both ANOVA approaches were p-value adjusted by a 2-stage step up Bejamini Krieger Yukatelli false discovery correction rate with Q set to 5%. Discovery is denoted as a p-value adjusted for multiple comparisons, q<0.05. Before statistical analyses were performed, data were searched for outliers and removed via Grubbs outlier testing (alpha=0.05). All statistical analyses were performed in GraphPad Prism (v10.4.1) with each statistical test described in figure legends.

## Acknowledgments

We thank the veterinary staff in the animal research facility for their outstanding animal care and Audrey L. Spiegelhoff for her technical assistance.

